# Fast and Robust Segmentation of Copy Number Profiles Using Multi-Scale Edge Detection

**DOI:** 10.1101/056705

**Authors:** Ivo W Kwee, Andrea Rinaldi, Cassio Polpo de Campos, Francesco Bertoni

**Affiliations:** Lymphoma and Genomics Research Program, Institute of Oncology Research, Bellinzona, Switzerland.; Dalle Molle Institute for Artificial Intelligence, Manno, Switzerland.; Swiss Institute of Bioinformatics, Lausanne, Switzerland.; Lymphoma Unit, IOSI, EOC, Bellinzona, Switzerland.

## Abstract

Raw copy number data is highly dimensional, noisy and can suffer from so-called genomic wave artifacts. We introduce a novel method based on multi-scale edge detection in derivative space. By using derivatives, the algorithm was very fast and robust against genomic waves. Our method compared very well to existing state-of-the-art segmentation methods and importantly outperformed these if noise and wave artifacts were well present.

## INTRODUCTION

Copy number aberrations (CNA) are a hallmark of cancer. With the coming of the current high density micro arrays, CNAs can be efficiently measured at several million locations in the genome. Segmentation of the copy number signal is a fundamental step before subsequent statistical analysis. Because of the high number of probes and because the aCGH data is very noisy, we need algorithms that are fast and robust.

A great number of segmentation algorithms have been proposed. A first class of methods employ some kind of local smoothing of the raw signal using, for example, moving average, wavelets (Ben-Yaakov and Eldar, 2008), or adaptive weights smoothing (Hupé 2004). With these methods, the data is not explicitly segmented but usually one proceeds by detecting contiguous altered segments by thresholding for loss and gain.

A special class of segmentation methods may be referred to as ‘piece-wise constant model’ (PCM) based methods that include CBS (Olshen *et al.*, 2004) and mBPCR (Rancoita *et al.*, 2009). These methods model the underlying true copy number profile as a piece-wise constant function. In theory, this is a reasonable assumption because the DNA breaks in segments and between two genomic breakpoints we expect the copy number to be constant. In practice, however, PCM-based segmentation is severely hampered by two problems: wave artifacts and hypersegmentation. ‘Genomic waves’ have been reported in the raw signal of genotyping arrays of both Affymetrix and lllumina (Nannya *et al.*, 2005; Komura *et al.*, 2006; Marioni *et al.*, 2007). These artifacts are exposed as local ‘wave-like fluctuations of the base line of different characteristic lengths from tens to hundreds of Mb. The cause of the genomic waves is not clear but seem to be related with the local GC content. The strength of the waves depends on the sample and may be related to the amount of starting material or differences in PCR conditions (Nannya *et al.,* 2005). These artifacts cause false positives and adversely limit the detection of (real) low contrast features, in particular for CNV detection. Furthermore, because of their pseudo-random nature, the artifacts seem dangerously ‘convincing’ as real copy number aberrations. A variety of methods for the correction of genomic wave artifacts have been proposed. These methods include regression on length and GC content (Nannya *et al.*, 2005; Leprêtre *et al.*, 2010), estimation and subtraction of the wave using lowess (Marioni *et al.*, 2007), and normalization using reference samples (van der Wiel *et al.*, 2009).

Finally, we define a last group of (copy number) segmentation methods that uses first derivative statistics to detect edges that correspond to chromosomal breakpoints. In image processing, edge detection has a long history and is one of the fundamental methods for image segmentation. The Canny edge detector, developed in 1986 by John F. Canny, is known to many as the optimal edge detector and is still one of the most competitive methods to date. Although edge-based segmentation for copy number profiling has not received much attention, we believe that edge-based segmentation for copy number data which can exhibit genomic waves and/or hypersegmentation has the better assumptions and less problems than the more established PCM based methods. To the best of our knowledge, FACADE is the only genomic segmentation method that uses edge detection in derivative space (Coe *et al.*, 2010). Compared to FACADE, our method uses multiple scales for better sensitivity and uses robust outlier statistics to determine edges, while the former detects edges at a single scale and fixes the number of edges.

We refer to our method as FFSEG (first-derivative based fast segmentation). Our proposed method is extremely fast and, more importantly, very robust against genomic waves artifacts and hypersegmentation. Briefly, FFSEG uses first derivatives of the raw signal and detects edges on multiple scales using robust outlier statistics. The use of the first derivative instead of the actual signal directly eliminates the problem of genomic waves, while robust outlier detection adapts to the noise of the individual sample and avoids hypersegmentation. The edge detection on multiple scales ensures good sensitivity of short high-contrast CNVs and at the same time broad low-contrast regions. The FFSEG code (in R) is available upon request.

## METHODS

Our starting point is the noisy, non-segmented, raw copy number signal that has been properly normalized and log2-transformed but not necessarily corrected for genomic waves. For a basic implementation, we defined a signed Gaussian edqe-detection (SGED) kernel as:

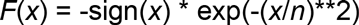

where *x* is the position index and *n* is a scale parameter. We preferred this choice of kernel rather than the more commonly used derivative of a Gaussian (Coe *et al.*, 2010) because it showed better localization of the edge positions due to the sharp boundary at x=0. For each chromosome, first derivatives were computed by convolving the raw signal with the SGED kernel. Edges were detected as statistically significant outliers, p<0.05 after Bonferroni multiple test correction. The detection of edges was repeated at multiple scales, n = (8, 16, 32, 64, 128, 256, 512, 1024, 2048). We found that this set of scales detected most edges in our test cases. The set of edges were then combined and duplicate positions were removed. This final set served as candidate positions of true chromosomal breakpoints. A subsequent pruning step removed breakpoint candidates by testing the mean signal of its flanking regions (t-test, p<0.05). Finally, we estimated the copy number level for each region by its mean value. This basic implementation worked well, however for larger scales the algorithm tended to become rather slow. To speed up, for each scale, we decided to firstly bin the raw signal onto a coarse resolution, detected the approximate positions of edge candidates and refined the edges by reestimating their positions with the SGED kernel using the full resolution of probes. This sped up the algorithm considerably without loss of accuracy.

**Figure 1.**
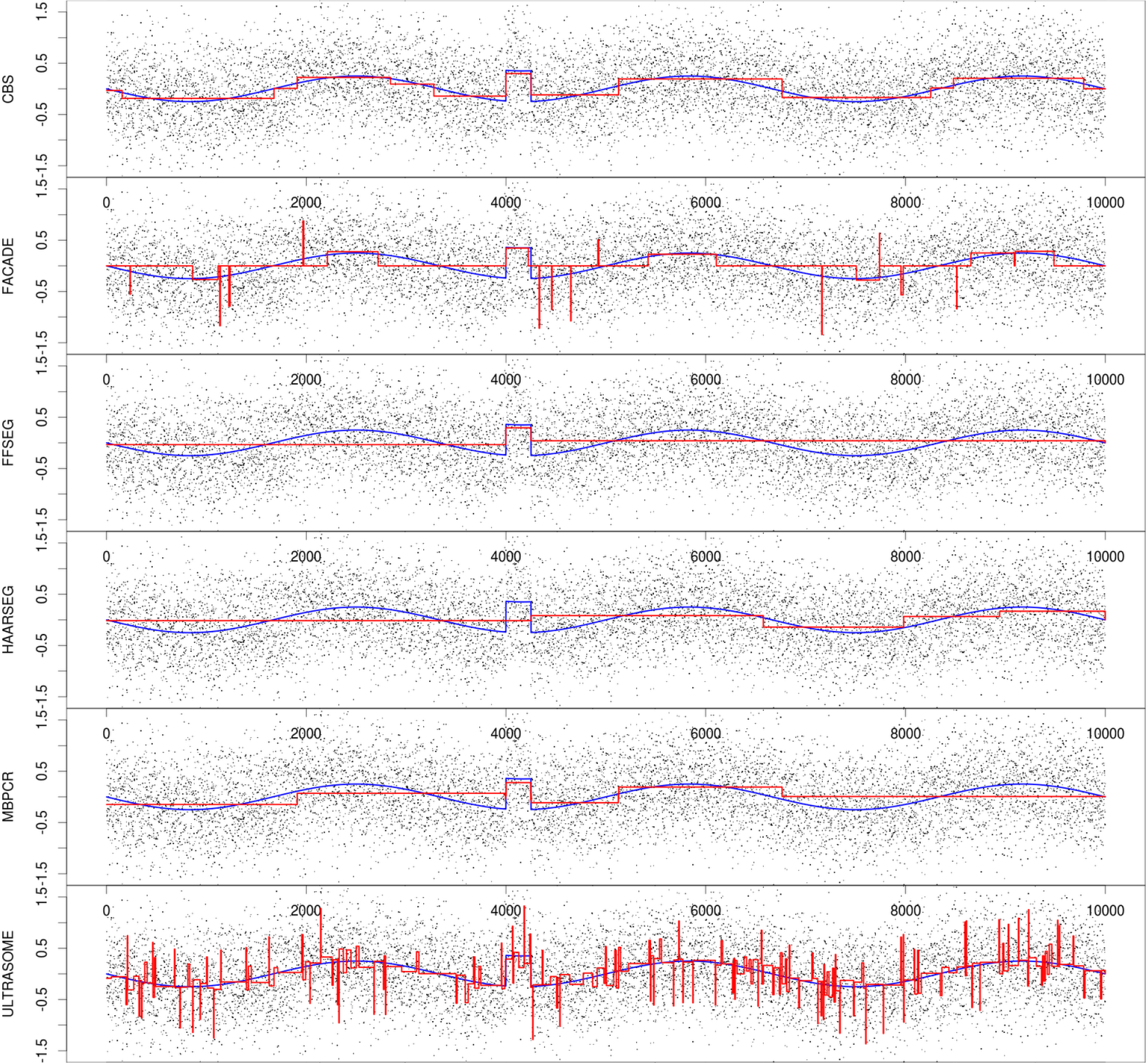
Comparison of several segmentation algorithms on an artificial example illustrating the effects of genomic waves. CBS, FACADE and mBPCR locate the aberration around x=4000 but are sensitive to the waves. Ultrasome (and to a lesser extent FACADE) suffers from spurious hypersegmentation, while HaarSeg fails to locate the aberration. FFSEG locates the aberration correctly and is robust against the genomic wave. The blue line indicates the “true” genomic profile showing a small copy number gain region embedded in the genomic wave. The black dots are raw copy number data. The red lines correspond to the segmentation results of the different algorithms, x-axis: genomic location, y-axis: log2ratio.

## RESULTS

*Simulated data*. To illustrate the features of FFSEG we created simulated data and compared its segmentation results to those using the state-of-the-art segmentation methods: CBS (Olshen 2004), mBPCR (Rancoita *et al.*, 2010), Ultrasome (Nilsson *et al.*, 2009), HaarSeg (Ben-Yaakov and Eldar, 2008) and the forementioned FACADE algorithm. Our simplified test case consisted of a small aberration embedded in a noisy wave pattern. We have chosen model parameters reflecting realistic values: ‘wave amplitude’=0.25, ‘noise SD’=0.60, ‘wave length’=3333, ‘number of probes’=10000, ‘aberration width’=250 probes, ‘aberration height’=0.60. Figure 1 shows the results of the different segmentation methods. If not otherwise mentioned, we used the default parameters for the different algorithms. We see that CBS correctly modeled the aberration segment but also closely modeled the waviness of the baseline, and it will be difficult to differentiate between the real aberration and the tops of the wave artifact that appear as ‘gains’. FACADE identified the aberration but suffered too from the genomics wave artifact and additionally showed spurious peaks. HaarSeg failed to localize the aberration. With more sensitive parameters, however, HaarSeg identified the gain but showed spurious noise artifacts elsewhere (results not shown). mBPCR correctly located the aberration but was still sensitive to the genomic wave, however less sensitive than CBS. The segmentation result of Ultrasome was extremely noisy and followed closely the genomic wave. Finally, the figure shows that only FFSEG correctly identified the aberration while completely ignoring the wave artifact of the baseline. Running times were 0.833 seconds for CBS, 1440 seconds for mBPCR, 0.315 seconds for UltraSome, 0.063 for HaarSeg and 0.015 seconds for FFSEG.

*Real data*. We have run FFSEG on real data. The example data (sample X5098) was taken from the public data set of tumor copy number profiles (van de Wiel *et al.*, 2009) and represents a sample with considerable genomic wave artifact. A set of normal reference sample were available. We compared FFSEG and CBS on the raw (non-corrected) data and the wave-corrected data using NoWaves (van de Wiel *et al.*, 2009). Note that for the wave-correction a set of normal references (either paired or not paired) must be available which is not always the case. In Figure 2, we can see that on non-corrected data CBS suffered considerable from wave artifacts and hypersegmentation (Figure 2a), while it is much improved on the wave-corrected data. On the other hand, FFSEG showed stable segmentation on both the non-corrected data and wave-corrected data.

**Figure 2.**
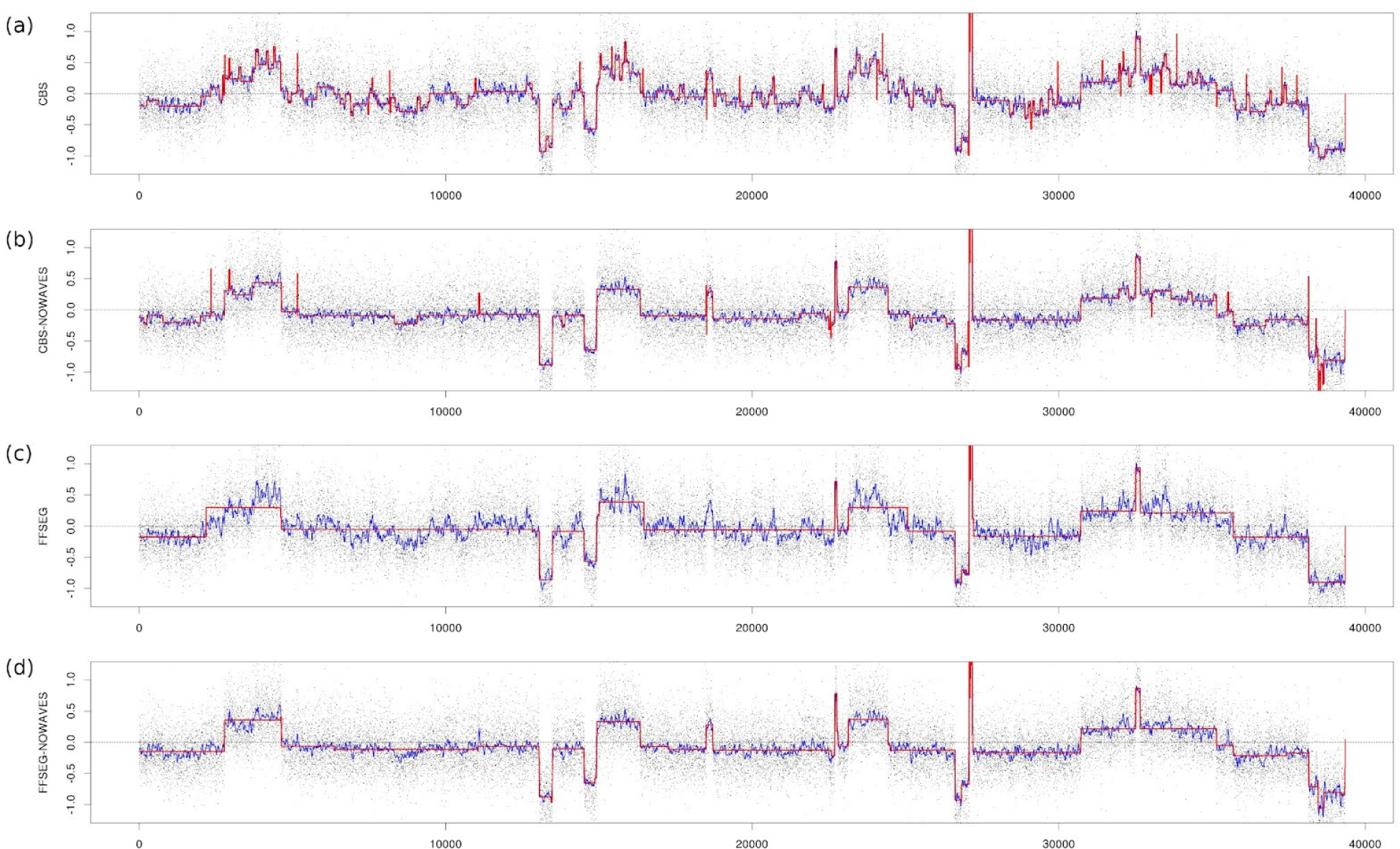
Comparison of FFSEG with CBS on real data with severe genomic wave artifacts. The data (sample X4861) is taken from the public data set of tumor copy number profiles (van de Wiel 2009). Using raw data FFSEG is robust against the genomic wave while CBS tends to hypersegmentation. Using NoWaves corrected data, FFSEG and CBS perform similarly but the FFSEG profile is still considerably cleaner, **(a)** CBS profile from raw data. Black dots denote raw data. Blue line denotes smooth data using moving average. Red line denotes segmentation, **(b)** CBS profile from NoWaves corrected data, **(c)** FFSEG profile from raw data, **(d)** FFSEG profile from NoWaves corrected data.

*Cytogenetic validation*. We validated FFSEG on available cytogenetic data of 396 cases of chronic lymphocytic leukemia (CLL) and compared its detection accuracy with that of CBS and mBPCR. The cytogenetic data represented lesions of different sizes: trisomy of chromosome 12, deletion of the small arm of chromosome 17, a region of loss at 11q (targeting *ATM*), and an interstitial loss at 13q around the *MIR15A/MIR16-1* cluster. Raw data was acquired using Affymetrix SNP 6.0 mapping array and normalized using the manufacturer provided Hapmap reference. The array has more that 1.8 million copy number probes. Segmentation of a single copy number profile, on an 8-core 2.0GHz Intel Linux workstation, took a few seconds with FFSEG, about 1 minute with CBS, and several hours with mBPCR. Figure 3 summarizes the results. FFSEG compares well with CBS and mBPCR for the larger lesions (trisomy 12, del17p, del11q) but was slightly more conservative for smaller deletions (del13q).

**Figure 3.**
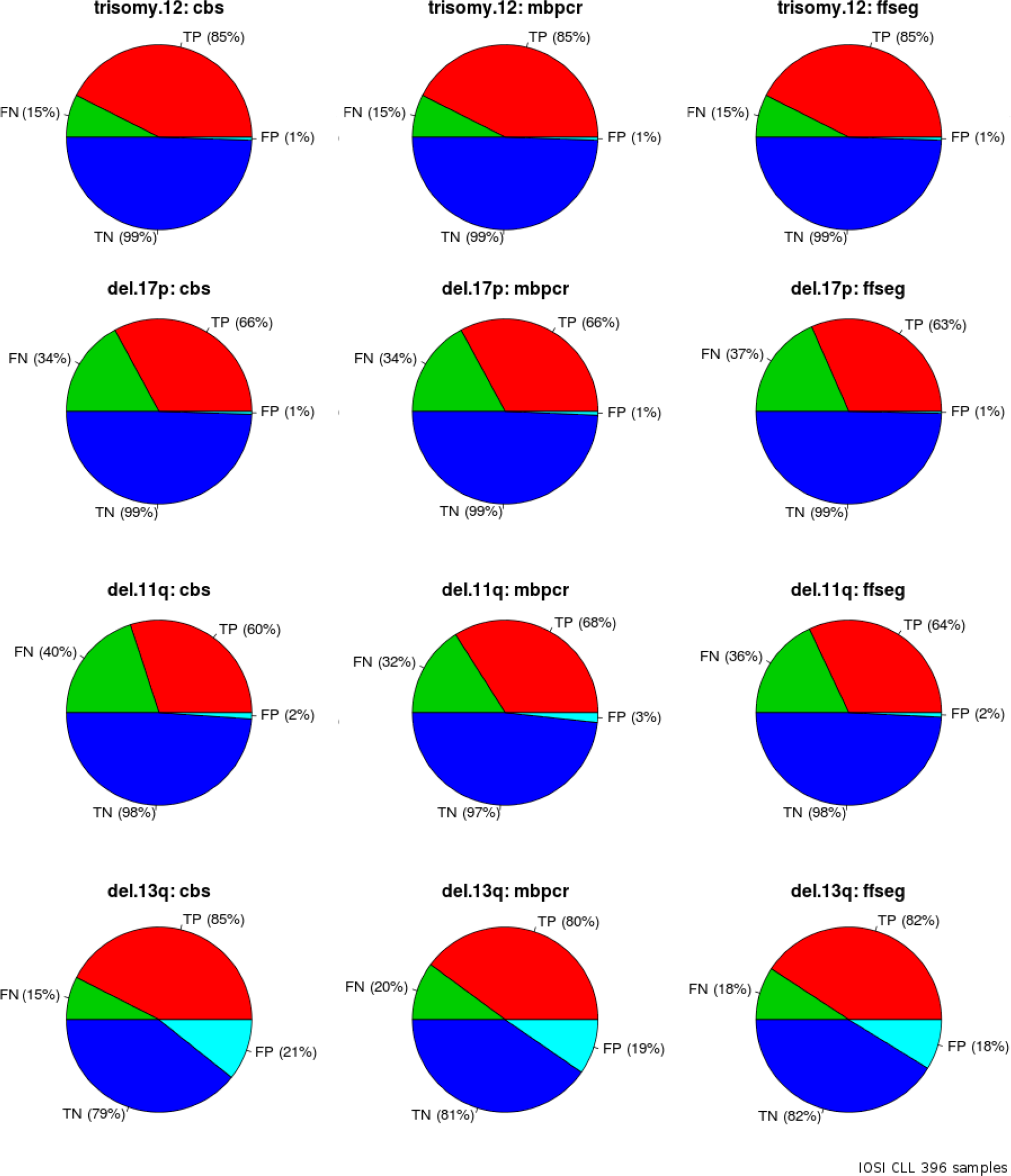
Detection accuracy of genomic lesions of CBS, mBPCR and FFSEG on cytogenetic validated data of 396 cases of chronic lymphocytic leukemia. For the larger lesions (trisomy12, del17p, del11q) FFSEG compares well with CBS and mBPCR. For the smaller lesion (del13q), FFSEG seems slightly more conservative than the other methods showing a lower false positivity (FP) and lower true positivity (TP).

## CONCLUSION

We have introduced a novel fast and robust algorithm (FFSEG) for the segmentation of copy number profiles that is based on multi-scale edge detection using derivatives. A drawback might be that for the derivative outlier detection FFSEG needs a good amount of probes to determine the null distribution, so the method might not be suitable for low density arrays. For relatively easy cases, FFSEG compared well with the current state-of-the-art piecewise-constant model based algorithms, and was the fastest method. More importantly, for more complex scenarios, FFSEG showed to be very robust against so-called genomic wave artifacts and hypersegmentation.

## ACKNOWLEDGMENTS

This research has been supported by grants from the Nelia and Amedeo Barletta Foundation, Helmut Horten Foundation and San Salvatore Foundation. The authors declare no conflict of interest.

## REFERENCES

Nannya, Y. et al. (2005). A robust algorithm for copy number detection using high-density oligonucleotide single nucleotide polymorphism genotyping arrays. Cancer Res., 65, 60716079.

Komura, D. (2006). Genome-wide detection of human copy number variations using high-density DNA oligonucleotide arrays. Genome Res. 16(12): 1575–1584.

Van de Wiel, M.A. etal. (2009). Smoothing waves in array CGH tumor profiles. Bioinformatics, 25, 1099–1104

Marioni, J.C. (2007). Breaking the waves: improved detection of copy number variation from microarray-based comparative genomic hybridization. Genome Biol., 8(10): R228.

Leprêtre, F. (2010). Waved aCGH: to smooth or not to smooth. Nucleic Acids Res. 38(7).

Canny, J. (1986). A computational approach to edge detection. IEEE Trans. Pattern Anal. Mach. Intell. 8, 6, 679–698.

Olshen, A.B. (2004). Circular binary segmentation for the analysis of array-based DNA copy number data. Biostatistics. 5(4):557–72.

Nilsson, B. et al. (2009). Ultrasome: efficient aberration caller for copy number studies of ultra-high resolution. Bioinformatics. 25 (8):1078–1079.

Rancoita, P.M.V. etal. (2009). Bayesian DNA copy number analysis. BMC Bioinformatics 10:10.

Ben-Yaacov, E. and Eldar, Y.C. (2008). A fast and flexible method for the segmentation of aCGH data. Bioinformatics, 24 (16):i139–i145.

Coe, B.P. et al. (2010). FACADE: a fast and sensitive algorithm for the segmentation and calling of high resolution array CGH data. Nucleic Acids Research, 2010, 1–7.

